# QuantUMS: uncertainty minimisation enables confident quantification in proteomics

**DOI:** 10.1101/2023.06.20.545604

**Authors:** Franziska Kistner, Justus L. Grossmann, Ludwig R. Sinn, Vadim Demichev

## Abstract

Mass spectrometry-based proteomics has been rapidly gaining traction as a powerful analytical method both in basic research and translation. While the problem of error control in peptide and protein identification has been addressed extensively, the quality of the resulting quantities remains challenging to evaluate. Here we introduce QuantUMS (<u>Quant</u>ification using an <u>U</u>ncertainty <u>M</u>inimising <u>S</u>olution), a machine learning-based method which minimises errors and eliminates bias in peptide and protein quantification by integrating multiple sources of quantitative information. In combination with data-independent acquisition proteomics, QuantUMS boosts accuracy and precision of quantities, as well as reports an uncertainty metric, enabling effective filtering of data for downstream analysis. The algorithm has linear complexity with respect to the number of mass spectrometry acquisitions in the experiment and is thus scalable to infinitely large proteomic experiments. For an easy implementation in a proteomics laboratory, we integrate QuantUMS in our automated DIA-NN software suite.

## Introduction

Liquid chromatography coupled to mass spectrometry (LC-MS)-based bottom-up proteomics is a highly dynamic field, with recent years being marked by rapid technology development, which is enabling novel applications and empowering existing ones. Faster and more sensitive instruments, novel acquisition methods and advanced data processing approaches have pushed the limits of throughput and sensitivity, with concomitant reduction of costs and advances in reproducibility. This has led to proteomics becoming a powerful and widely used tool for both discovery of fundamental biology and translational applications, as well as one of potential pillars of personalised medicine^1^.

Data-independent acquisition (DIA)^2–4^ proteomics has become highly popular in recent years, as the instrumentation and analysis software advances have addressed most of its drawbacks, while strengthening such advantages of DIA as high proteomic depth and data completeness along with improved quantitative performance^1^. Further, a number of recent DIA technologies also inherently manifest what has been the main benefit of data-dependent acquisition (DDA), a reliable fragment to precursor mass assignment, promising to expand the applicability of DIA even further^5–8^. A range of transformative improvements in the sensitivity of DIA^9–15^ combined with multiplexed DIA^16–22^ further promise an expansion of DIA towards applications that have previously relied on targeted methods such as selected or parallel reaction monitoring.

Much of the progress in DIA proteomics has been driven by data processing software improvements^23–30^, with novel algorithms yielding rapid gains in proteomic depth. Advanced machine learning approaches now enable confident matching of peptides to recorded signals even if the latter are highly noisy and are affected by signal interferences from co-eluting and co-fragmenting peptide species. This has raised the question of how quantitative these extra identifications are, and to what extent they benefit the biological inference from the experiment. This issue is particularly important given recent interest in workflows that stretch the capabilities of the instruments and generate challenging to process data by boosting chromatography throughput up to hundreds of samples per day, including for high-sensitivity applications, such as single-cell proteomics or spatial tissue proteomics^5,9,11,21,31–37^.

Significant effort has been directed towards developing computational methods that can improve quantitative reproducibility, precision and accuracy of proteomic experiments. These include deconvolution of spectra^38^, selection of peptide fragment ions based on the signal quality^21,26,39^, as well as protein quantification through aggregation of multiple parallel sources of quantitative information, such as peptide-level MaxLFQ for DDA^40^ and fragment-level MaxLFQ for DIA^41^ or directLFQ^42^. Advanced methods for error control and missing data handling have also been developed for statistical analysis of proteomics data, as discussed and benchmarked recently^43,44^. However, while peptide identification error rates are well controlled by statistically-justified target-decoy competition methods, quantification errors are currently impossible to estimate.

Here we introduce QuantUMS (<u>Quant</u>ification using an <u>U</u>ncertainty <u>M</u>inimising <u>S</u>olution), a novel concept for accurate and reliable quantification in proteomics. QuantUMS draws upon the quality information available for individual signals recorded by the mass spectrometer. This allows QuantUMS to integrate multiple signals associated with each precursor and protein in a statistically-justified way by propagating the quantification uncertainty estimates that it obtains from the quality information. Furthermore, QuantUMS compares multiple parallel sources of quantitative information present in the data and thus obtains concordance metrics that allow it to leverage machine learning to optimise its hyperparameters. In addition, QuantUMS generates quantification accuracy metrics for individual precursor and protein quantities, allowing for effective filtering of the processed data for better downstream statistical inference. We implement QuantUMS as a module in our automated and easy to use DIA-NN software suite ^26^ and show that, when applied to DIA data, QuantUMS boosts quantitative precision and enhances differential expression analyses. We also demonstrate that QuantUMS is capable of eliminating the ratio compression bias that so far has been inherent to untargeted proteomics data.

## Results

### Principle of QuantUMS

In each acquisition, the mass spectrometer records multiple signals for each detected peptide precursor: the signal from the unfragmented precursor in MS1 mode, as well as the signal from each of its fragment ions in MS/MS mode. Integrating all of these signals should naturally allow to achieve higher accuracy and precision than using just one of them. However, all of these measurements are subject to errors caused by random noise^45^ and ‘interfering’ signals from precursors with the same mass (MS1) or from precursors that share some of the fragment ions and are co-isolated for fragmentation^4^. Ideally, the integration method needs to leverage some uncertainty measures, calculated for each signal.

Previously, we devised and implemented in our DIA-NN software a straightforward but highly effective approach for quantification from DIA data, wherein several fragment ions are selected for each precursor, in a cross-run manner, and then the sum of their respective signals in each acquisition is used to represent the respective precursor quantity. The selection was performed by choosing the top three fragments with highest average correlation scores across the experiment, wherein the correlation score of each fragment is calculated by DIA-NN as part of the precursor identification process for a particular acquisition and represents how well its extracted elution profile is aligned with extracted elution profiles of other fragments of the same precursor. This strategy enabled filtering out signals which are strongly affected by interferences in multiple runs, thus greatly boosting the quality of quantification^26^. However, this approach based on quality scores averaged across runs is still subject to fold-change interference-related errors in individual acquisitions. Further, it has a significant drawback of discarding much of the information encoded in the raw data, such as the measured MS1 signal. In fact, in our later work on multiplexed DIA proteomics for low sample amounts^21^, we quantified precursors at the MS1 level instead, as this led to higher quantification accuracy on an Orbitrap setup that used 50% MS1 duty cycle. Others have also observed that leveraging both the MS1 and MS2 precursor quantities empowers downstream statistical analysis^46^.

We therefore aimed to develop a quantification method that would draw upon all the information on the precursor that was recorded by the mass spectrometer, as well as to integrate this information guided by the quality scores available for individual recorded signals, in a statistically-justified way. Below we outline the concept behind QuantUMS, with a detailed description of the algorithm provided in Methods.

QuantUMS (Figure 1a) takes as input the set of signal intensities corresponding to the quantitative features – precursor itself recorded at the MS1 level as well as its fragment ions recorded at the MS/MS level – for each peptide precursor in each acquisition where it was identified, as well as a single quality score per signal. While in principle different kinds of quality scores can be used, the QuantUMS implementation in DIA-NN that we describe here (Methods) leverages the correlation-based similarity score that DIA-NN calculates when comparing an extracted ion chromatogram of a feature with that of the selected ‘best’ fragment ion it defines for each putative elution peak^26^. QuantUMS then models the log-space variance of the measured signal (‘log-variance’), a proxy for quantification uncertainty, as a function, defined by the algorithm hyperparameters, of the signal intensity and its quality score. These log-variance estimates for individual feature signals are then used to construct a formula for the quantity of each precursor identification that minimises its estimated log-variance.

**Figure 1.**
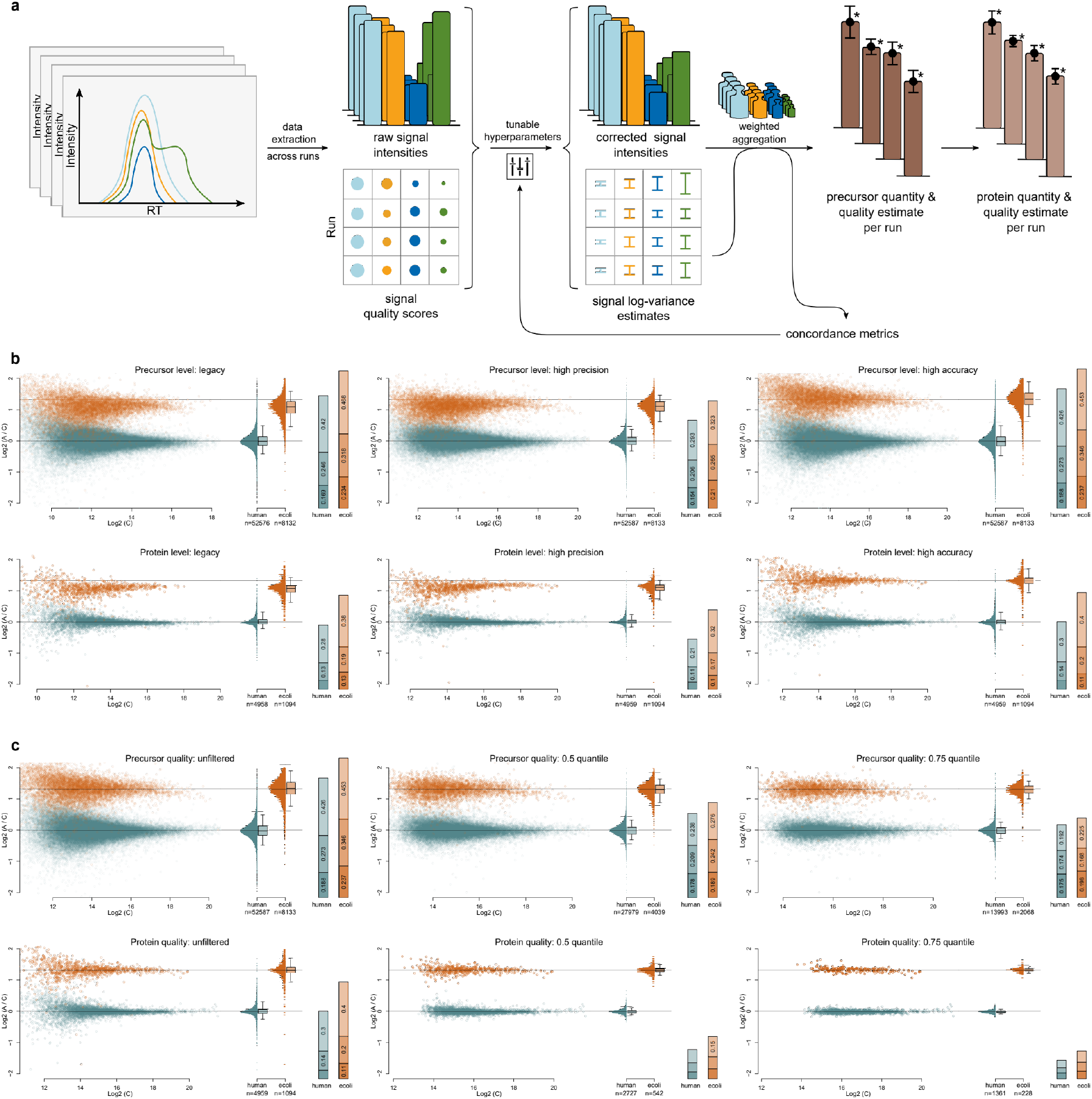
QuantUMS performs statistically-justified minimisation of quantification uncertainty. **a**, QuantUMS takes as input integrated signals of individual quantitative features (MS1 precursor and MS/MS fragment ions), as well as their quality scores. These allow QuantUMS to correct signal intensities to remove bias and to estimate their log-variances, in a process controlled by hyperparameters. The signal log-variance estimates cater for a statistically-justified weighted aggregation of signals, to obtain precursor quantities, and can also be propagated through weighted aggregation, to obtain log-variance estimates for these precursor quantities, allowing QuantUMS to report a ‘quantity quality’ metric for each precursor. QuantUMS also establishes concordance metrics based on comparing parallel channels of quantitative information, enabling it to optimise the hyperparameters using machine learning. Precursor quantities and their log-variance estimates are likewise aggregated to obtain protein quantities and the respective quantity quality metrics for proteins. **b**, Three mixtures A, B and C of human (K562) and *E*.*coli* tryptic digests with proportions A:B:C being 1:1:1 (human) and 50:33:20 (*E*.*coli*) were recorded^37^ using a 5-minute analytical flow gradient on timsTOF Pro and analysed using legacy and QuantUMS quantification methods. Resulting precursor and protein ratios between mixtures A and C are visualised. On the boxplots, boxes correspond to the interquartile range with the median indicated, whiskers extend to 5%-95% quantiles. Horizontal lines indicate the expected ratios for each species digest. Stacked bar plots indicate log2-standard deviations of A:C ratios within three intensity bins, with the lowest intensity bin at the top. **c**, The effect of run-specific and precursor-specific or protein-specific, respectively, quality filtering enabled by QuantUMS, data shown for the high-accuracy mode.

Essential to QuantUMS is the idea that different quantitative features each can produce an estimate of a quantity of a precursor in a particular acquisition, relative to other acquisitions. These estimates can be compared between features, and high concordance between these estimates is likely to be indicative of their accuracy. QuantUMS uses machine learning to infer hyperparameters that maximise concordance between precursor quantity estimates obtained using different features.

Quantitative proteomics data is subject to signal interferences, which tend to cause ratio compression and hence biased quantification for precursors that are not highly abundant. In this work we present a solution to this problem: we introduce a mechanism in QuantUMS, which is capable of effectively eliminating such bias, at the cost of a limited loss of precision. QuantUMS incorporates a bias correction in its calculations, based on signal quality scores and intensities, which is controlled by hyperparameters. To tune these, we devise a way to empirically assess and minimise the degree of bias manifested by interference-affected signals, without requiring any knowledge of the experiment design. The QuantUMS module in DIA-NN implements a configurable parameter which allows to control the impact of this mechanism and thus balance precision and accuracy, as well as implements two pre-configured modes termed high-precision and high-accuracy, which we benchmark in the present work.

Controlling for quantification errors has so far been challenging in proteomics. Indeed, the only widely used solution has been to filter the dataset based on precursor or protein coefficients of variation (CVs). This approach does not control for interference-caused errors that might severely impact the accuracy of quantification while preserving precision, nor does it account for errors that only manifest in some of the acquisitions and thus do not have a significant negative impact on the CV values. This necessitates the need for laborious, potentially biased and sometimes technically impossible manual checks of extracted ion chromatograms for each acquisition and peptide of interest, to ensure confidence of observations pertaining to specific proteins. With QuantUMS, we address this problem by introducing a precursor- and acquisition-specific quantity quality metric, which, while not completely eliminating the need for manual checks aimed at maximum confidence, significantly reduces it in many applications.

In QuantUMS we devised an algorithm that ensures confidence in individual precursor quantities. The procedure applied by QuantUMS is based on combining multiple signals in a way to minimise the resulting log-variance, which can then be estimated as a function of the algorithm hyperparameters. QuantUMS then calculates the bias of these log-variance estimates, that is, determines if they are too conservative or too optimistic, by comparing the observed deviations between precursor quantity estimates obtained using different quantitative features with the expected log-variances of these quantity estimates. It then reports a ‘quantity quality’ score, for each precursor identification, based on precursor quantity log-variance estimates calibrated to remove the bias.

We further extend QuantUMS with a protein quantification module, which likewise relies on quality metrics to weigh multiple channels of quantitative information and likewise reports the resulting protein quantity quality metric (Methods).

### QuantUMS improves precision and boosts accuracy by eliminating bias

First, we benchmarked QuantUMS on an LFQbench-type^47^ dataset that we had previously acquired using a 5-minute gradient on an analytical flow liquid chromatography system coupled to timsTOF Pro operated in dia-PASEF mode^37^. In this experiment, wherein human (K562) and *E*.*coli* tryptic digests were mixed in three different proportions A, B and C (A:B:C ratios 1:1:1 for human and 50:33:20 for *E*.*coli*, respectively), the ability of the instrument and the data processing software to accurately and precisely recover those proportions as well as detect the differential levels of *E*.*coli* peptides and proteins is evaluated. We independently verified the accuracy of *E*.*coli* sample dilution used to prepare mixtures A and C by analysing the precursor identifications with high quality MS1 signals (Supplementary Figure S1). We compared the high-precision (default) and high-accuracy modes of QuantUMS to the ‘legacy’ DIA-NN mode (Figure 1b), which represents the default quantification strategy employed by the previous versions of DIA-NN^26^. We observed that the high-precision mode of QuantUMS boosted the precision of inferred ratios between mixtures, yielding a substantial improvement of log2-standard deviations of precursor and protein quantity ratios, with the greatest effect seen for the low intensity precursors (over 1.4-times improvement) and proteins (1.3-times, human, and 1.2-times, *E*.*coli*). The high-accuracy mode of QuantUMS did not result in a significant precision improvement over the legacy DIA-NN mode, however the median inferred ratios in this mode matched the expected ones, thus eliminating the ratio compression bias that is observed in legacy and high-precision modes.

We further examined the effect of filtering the dataset based on the acquisition-specific precursor- and protein-level quantity quality metrics introduced in QuantUMS (Figure 1c). As expected, we observed that such filtering tends to retain only accurately quantified precursors and proteins, with 0.75 quality quantile protein-level filtering resulting in almost perfect protein ratios (Figure 1c, bottom right panel).

Although QuantUMS contains a hyperparameter tuning step, it can also be instructed to use a particular set of hyperparameters, potentially trained on a different data set. Using the same experiment, we examined the effect of the training data set on QuantUMS performance in the high-accuracy mode, by optimising the hyperparameters only on samples A+C, A or C separately, and then applying them to the whole data set A+B+C (Supplementary Figure S2). In each scenario, we observed a performance similar to the one demonstrated when training hyperparameters on A+B+C (Figure 1b, rightmost panels). This indicates that the optimised hyperparameters of QuantUMS reflect the inherent properties of the LC-MS setup and the sample matrix, but do not reflect the experiment design.

We investigated what effect the enhanced performance of QuantUMS has on differential expression analyses. First, we examined the same mixed species dataset, in which the ground truth is known and can be used to determine the effective false discovery rate (FDR). Specifically, we applied Welch t-test to each precursor or protein, and plotted the numbers of true hits (*E*.*coli*) against the FDR represented by the ratio of human to *E*.*coli* hits (Figure 2a). Here we consciously went for the simplest statistical test, as our goal is to benchmark the quality of precursor and protein quantities rather than the statistical approach.

**Figure 2.**
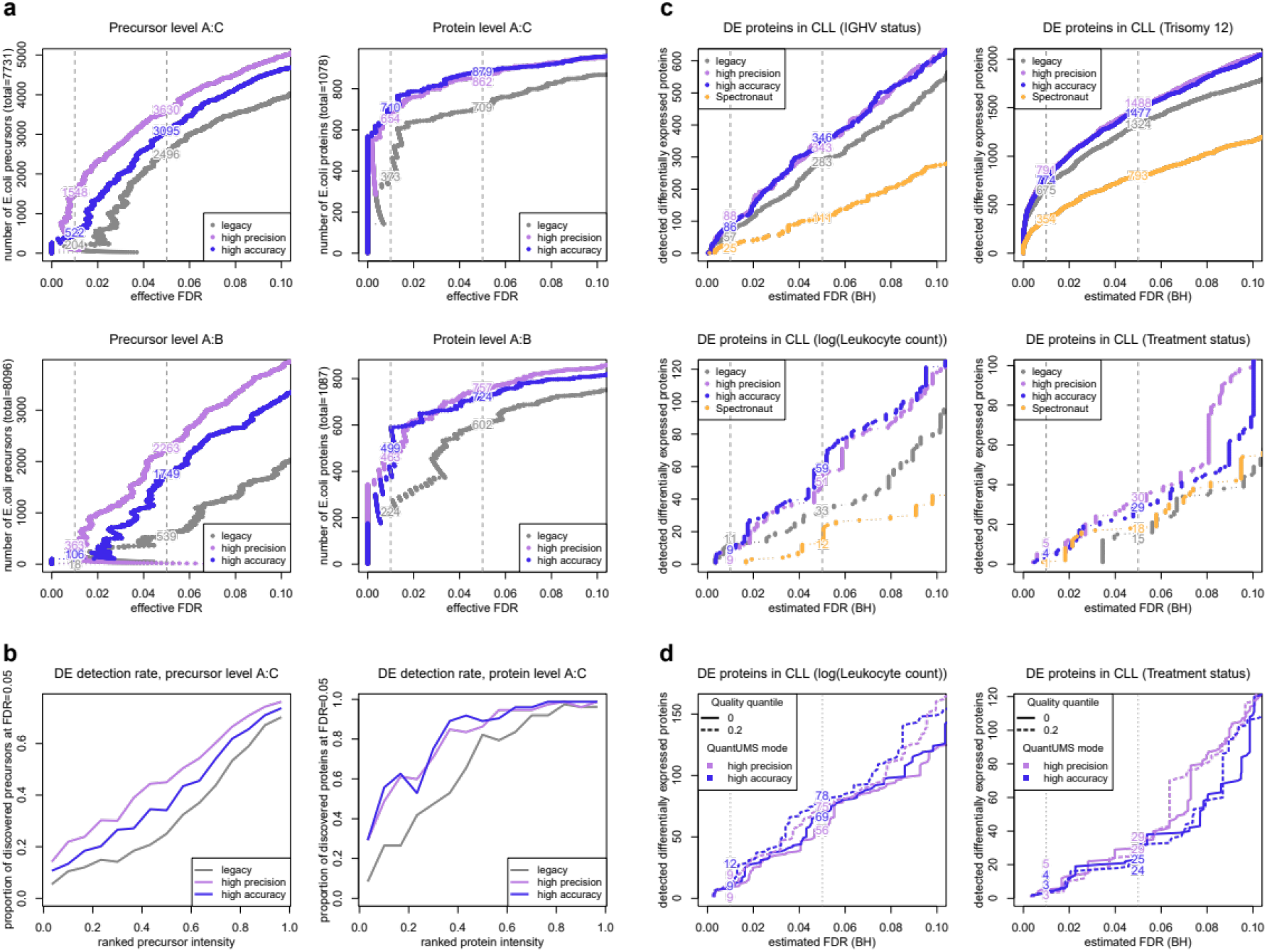
QuantUMS boosts the sensitivity of differential expression analysis. Three mixtures A, B and C of human (K562) and *E*.*coli* tryptic digests with proportions A:B:C being 1:1:1 (human) and 50:33:20 (*E*.*coli*) were recorded^37^ using a 5-minute analytical flow gradient on timsTOF Pro and analysed using legacy and QuantUMS quantification methods. **a**, Numbers of differentially expressed precursors and proteins plotted against the effective false discovery rate (true hits: *E*.*coli*, false hits: human) for A:C and A:B comparisons. **b**, Differential expression detection rate depending on the intensity rank for the *E*.*coli* 50:20 comparison (Methods). **c**, A comparison between legacy and QuantUMS analyses using DIA-NN, as well as to the originally published^48^ analysis using Spectronaut, on a dataset of 50 chronic lymphocytic leukaemia (CLL) samples recorded using a 100-min nanoLC gradient on timsTOF Pro. Numbers of differentially expressed proteins when testing against four phenotypic characteristics (limma, Methods) are shown. **d**, The effect of filtering the protein list using the QuantUMS protein quantity quality metric averaged across acquisitions on the number of differentially expressed proteins at a given FDR in selected tests.

Both QuantUMS methods performed superior to the legacy method, with the high-precision mode showing somewhat better performance than high-accuracy. We also observed greater advantage of QuantUMS at the precursor level than the protein level. This is in line with expectations, as protein quantities, obtained with either method, integrate multiple precursor quantities and are, therefore, inherently more precise. Thus, it is less challenging for the method to detect differential expression of proteins. Indeed, we see that the QuantUMS high-precision mode reports 871 differentially expressed proteins at 5% FDR for the A:C comparison, which is close to the total number of *E*.*coli* proteins (1078) detected in at least two replicates in both A and C. Likewise, the advantage of QuantUMS methods over the legacy method appeared even more significant for the A:B comparison, which features a tighter (50:33 as opposed to 50:20) ratio between the *E*.*coli* fractions and is thus more demanding with regards to quantitative precision. In either case, the relative advantage of QuantUMS was greater at higher confidence levels. Examining the rate of differential expression detection depending on the respective intensity rank, we see that the advantage of QuantUMS over the legacy method originates in its better ability to detect differential expression of low-abundant and hence more challenging to quantify precursors and proteins (Figure 2b).

To evaluate the performance of QuantUMS in an experiment that is also subject to biological variation and variation due to sample preparation in addition to quantitative variation introduced by the LC-MS, and thus should benefit less from the advantages of QuantUMS than a synthetic LFQbench-type benchmark, we re-processed a data set of 50 chronic lymphocytic leukaemia (CLL) samples acquired with a 100-minute nanoLC gradient on timsTOF Pro using dia-PASEF, which was analysed with Spectronaut in the original publication^48^. Testing for differential expression with respect to the phenotypic information available (Figure 2c; Methods), we observed greater numbers of proteins identified as differentially expressed by the QuantUMS methods in comparison to the legacy DIA-NN quantification, which in turn outperformed the original analysis with Spectronaut. The advantage of QuantUMS over legacy quantification was most prominent in case of the more challenging tests against leukocyte count and treatment status. We further observed that filtering the protein lists based on the averaged across-runs protein quantity quality metric is capable of improving the numbers of significant proteins at a given FDR in those tests (Figure 2d), by reducing the number of proteins considered and hence reducing the number of statistical tests to which Benjamini-Hochberg multiple testing correction is applied.

## Discussion

With QuantUMS, we address a long-standing problem of untargeted proteomics, that is the lack of quality control for peptide and protein quantities obtained in an experiment. We show that taking into account the quality information available for individual signals recorded by the mass spectrometer not only allows to improve quantitative performance per se, but to also produce effective quality metrics to ensure confidence in the data and further empower the subsequent statistical analysis.

So far, we have benchmarked the new method on DIA proteomics data, with the quality scores for recorded signals being calculated by our DIA-NN software. As opposed to relying on hand-picked thresholds like many legacy methods^26^, which usually can only be optimal for a narrow range of data sets, QuantUMS is driven by automatic optimisation using machine learning, and is hence highly flexible in terms of the input information it can accept. We therefore envision a potential for further improvements of QuantUMS through the introduction of multiple thorough signal quality metrics.

Furthermore, other kinds of proteomic acquisition approaches likewise generate multiple channels of quantitative information that can be leveraged by QuantUMS. These include selected and parallel reaction monitoring as well as other experiments that involve recording multiplexed MS/MS spectra, and application of QuantUMS to these can be explored in a follow-up work. We further envision significant potential for future improvements in quantitative proteomics to be achieved by integrating QuantUMS with downstream statistical analysis approaches, such as MSStats^43^ or Triqler^44^, to enable biological inference that is fully aware of all kinds of uncertainty, missingness and normalisation issues in the raw proteomics data.

Proteomics has a potential to significantly enhance personalised medicine, both for biomarker discovery and clinical decision-making. At the same time, reliable quantitative quality control is of particular importance for clinical applications^49–52^, while manual validation of LC-MS data in this setting might not be technically feasible and poses a risk of introducing a human bias. Automated peptide- and protein-level quality control, as enabled by QuantUMS, presents, therefore, a step towards reliable clinical proteomics.

## Methods

### QuantUMS algorithm

We first outline the core ideas behind the method and then discuss the enhancements implemented in QuantUMS. The QuantUMS algorithm is based on estimating the log-variance and log-bias of the measured signal intensity of a quantitative feature (either the MS1 precursor signal or an MS/MS fragment ion signal) using its value and a quality score. Specifically, given the integrated signal intensity S and a quality score C, it calculates transformed values 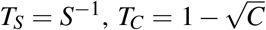 and 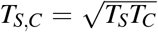. The implementation of QuantUMS in DIA-NN obtains the score *C* as the product of Pearson correlation and cosine similarity measures between the extracted elution profile of the feature in question and the smoothed elution profile of the fragment DIA-NN has labelled as ‘best’ during the identification process^26^. QuantUMS then models the log-variance of the signal as the linear combination of these transformed values, with coefficients corresponding to trainable hyperparameters. To enforce non-negativity of the coefficients, they are represented as squared values of hyperparameters. Likewise, QuantUMS models the log-bias as the linear combination of the square roots of the transformed values, and corrects signal intensities of features for their estimated bias when calculating precursor quantities. In total, this yields six hyperparameters.

QuantUMS iteratively optimises the precursor quantities, starting with the preliminary quantities calculated by summing selected fragment signals, as described for DIA-NN previously^26^. For a given precursor and an acquisition *i*, let *q*_*i*_ be the precursor log-quantity estimate obtained at the previous iteration and *s* _*f, i*_ – the logarithm of the bias-corrected signal intensity of quantitative feature *f*, in acquisition *i*. Then *x* _*f, j,i*_ = *q*_*i*_ + *s* _*f, j*_ − *s* _*f, i*_ estimates the log-quantity of the precursor in any acquisition *j*. To obtain the next-iteration estimate *x* _*j*_ of the precursor log-quantity in acquisition *j*, QuantUMS first takes a weighted average of *x* _*f, j,i*_ across acquisitions *i*, obtaining estimates *x* _*f, j*_ based on individual quantitative features, and then across features *f*, obtaining a single estimate. The principle of aggregation via weighted averaging implemented in QuantUMS is derived from the following statistical property. Given a set of uncorrelated random variables *Z*_*i*_ with equal means *μ* and variances var_*i*_, the estimate 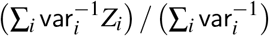 has the mean *μ* and the minimal, across all such linear combinations, variance equal to 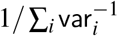, thus yielding the optimal estimate for *μ*. QuantUMS applies the same formula when it needs to aggregate multiple quantities given their estimated variances, thus obtaining both an estimate of their mean and the estimated variance of this aggregated value.

The above quantification method effectively implements a particular formula, which calculates the quantity of a precursor in each acquisition, based on the respective feature signal measurements in all acquisitions and their quality scores, as well as precursor quantity estimates obtained at the previous iteration. While this calculation is statistically justified given the assumptions of variances being known and of measurement errors being independent, variances cannot be precisely estimated in practice and the independence assumption likely does not hold strictly. We alleviate this drawback by having the formula depend on hyperparameters that are then tuned by machine learning. The idea here is that since each feature produces, for each acquisition, an estimate of the precursor quantity, the deviations between these estimates corresponding to different features are indicative of how accurate the quantity estimates are. QuantUMS hence tunes hyperparameters to minimise the empirically measured differences between quantity estimates obtained using different features.

This idea would perform poorly in practice, if the loss function for hyperparameter optimisation were calculated using all precursor identifications and all feature pairs without regard for their quality scores, and QuantUMS hence incorporates a more complex implementation. Specifically, for each precursor, QuantUMS selects a quantitative feature with the best average quality score across acquisitions in which the precursor has been identified confidently. Choosing this feature only across fragment ions or always selecting MS1 signal makes little difference in our benchmarks, so for better applicability of the method in case the experiment has poor quality MS1 information the selected feature *f*_*sel*_ is currently chosen only among fragments. QuantUMS then selects acquisitions in which this feature has a quality score above a certain margin (0.9 in the current implementation), that is, in these acquisitions it is likely to be almost free from signal noise or interference. The aggregation procedure via weighted averaging is then repeated only considering the selected acquisitions and considering the respective signal intensity of the selected feature instead of the previous-iteration quantity estimates *q*_*i*_, thus aggregating 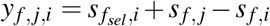 and yielding *y* _*f, j*_ estimates of 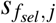. The first loss metric *l*_*prec*_ for hyperparameter optimisation, which drives optimisation towards minimising variance and hence maximising precision, is then calculated as the average of 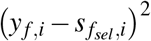 across acquisitions *i*, features *f* and all precursors for which such calculation is possible given the quality filtering margins. Another loss metric *l*_*acc*_ is designed to eliminate bias. Weighted average *μ*_*sel*_ and variance var_*sel*_ are calculated for the 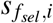 values considered across all precursors, using weights 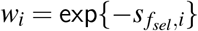, that is *w*_*i*_ is the inverse of the measured signal for the selected feature. QuantUMS then defines

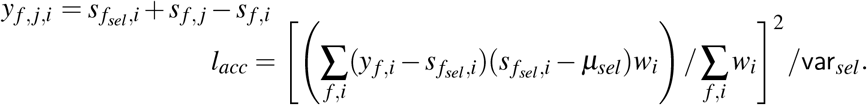

The intuition behind this formula is that ratio compression due to interfering signals occurs for low-abundant precursors. Minimising the *l*_*acc*_ loss therefore decorrelates the observed bias 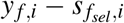 from 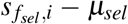, which serves as a proxy for precursor abundance. The final concordance metric is calculated as a linear combination of *l*_*prec*_ and *l*_*acc*_, with coefficients based on a user-defined balance between precision and accuracy. The pre-configured high-precision and high-accuracy modes of the QuantUMS module in DIA-NN give 50% and 10% weight to precision, respectively.

To enable hyperparameter optimisation, QuantUMS implements automatic differentiation. That is, all steps of aggregation via weighted averaging as well as the loss calculation are represented as differentiable arithmetic operations on hyperparameter vectors, and each calculation produces not only the output value but also the propagated gradient with respect to the hyperparameters. The latter can hence be optimised towards loss minimisation via a gradient descent. For QuantUMS it is essential to guarantee that optimised hyperparameters are better than their initial values, and thus the standard gradient descent procedure, which can lead to loss value spikes during optimisation, is not suitable. We therefore opted for backtracking gradient descent with Armijo line-search, which guarantees strict decrease of the loss function during optimisation, with a backtracking version of momentum applied^53^.

The algorithms described above produce not only quantity estimates but also log-variance estimates for each precursor identification. Given that assumptions of the statistical model used by QuantUMS are idealised and are not fully met by the empirical data, QuantUMS calibrates these log-variances by comparing the empirically calculated ∑_*i*_(*x*_*MS*1,*i*_ − *x*_*MS*2,*i*_)^2^ with its mathematical expectation based on the estimated variances of *x*_*MS*1,*i*_ and *x*_*MS*2,*i*_, where *x*_*MS*2,*i*_ is the quantity aggregated from *x* _*f, i*_ across all fragment features *f*. Given the calibrated log-variance estimate cal.var_*i*_, QuantUMS reports for the respective precursor the quantity quality metric as 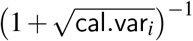 for acquisition *i*.

QuantUMS implements a range of enhancements to the core algorithms described above. First, we found the benefit of applying asymmetric quality filters to precursor identifications in runs *i* and *j* when calculating *x* _*f, j,i*_ = *q*_*i*_ + *s* _*f, j*_− *s* _*f, i*_. Specifically, both the average quality score and the quality score in acquisition *i* must be above certain margins for the feature *f*, for it to be used in this calculation. In contrast, applying a similar filter in acquisition *j* too appears detrimental. Enabling this asymmetry in quality filtering is one of the reasons for opting for the weighted averaging procedure as described for QuantUMS, as opposed to using weighted MaxLFQ. The other reason is the quadratic or cubic, depending on the implementation, computational complexity of MaxLFQ^40^ with respect to the number of acquisitions and the resulting challenges in scaling precursor-level MaxLFQ quantification to large experiments, as opposed to the linear complexity of QuantUMS. Second, quantities produced by QuantUMS are obtained via weighted averaging and QuantUMS also calculates the expected log-variances of these. Thus, QuantUMS also takes into account the variance of *q*_*i*_ in the formula *x* _*f, j,i*_ = *q*_*i*_ + *s* _*f, j*_ − *s* _*f, i*_, when propagating variance estimates through weighted averaging. Furthermore, the calculation of the variance of *s* _*f, i*_ in the above formula also incorporates *T*_*C*_ (defined above) that is multiplied by a tunable coefficient, adding an extra hyperparameter, where *C* is the average quality score for feature *f* across all acquisitions. In addition, QuantUMS adds an extra small value to all estimated variances for regularisation. It further adds log(10) · PEP to the estimated log-variance of a feature signal, where PEP is the posterior error probability of the respective precursor identification, ensuring that precursors identified with marginal identification confidence cannot be reported as having high quantitative confidence. Finally, QuantUMS also tracks how significantly the averaged expected variance of *y* _*f, i*_ deviates from the averaged empirically estimated one, by comparing *y* _*f, i*_ to 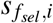, and adds extra regularisation to the loss function that places bounds on this deviation.

### Protein quantification in QuantUMS

We also incorporate a protein quantification algorithm in QuantUMS, derived from the MaxLFQ idea^40^, but implemented in a way that is variance-aware and enables protein quantity quality control. In MaxLFQ, protein quantities are found as a solution of a least squares problem that minimises the deviations between protein quantity log-ratios between acquisitions and empirical estimates of these log-ratios, obtained by taking the median of log-ratios between precursors mutually identified in the respective pair of acquisitions. QuantUMS here takes advantage of having estimated log-variance of each precursor quantity. Hence, a weighted median of precursor log-ratios is taken instead of a regular median, when estimating the log-ratio of protein quantities between two acquisitions. The respective least squares summand for the pair of acquisitions *i* and *j* in the MaxLFQ procedure is then assigned a weight equal to (∑_*k*_ var_*k,i*_ + var_*k, j*_)−1, where var_*k,i*_ is the estimated log-variance for the quantity of precursor *k* in acquisition *i*, thus giving higher weights to high quality precursor quantity ratios. Finally, QuantUMS also produces a quality metric for protein quantities, calculated as 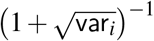, where 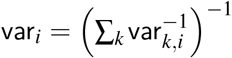. We note that protein quantification in bottom-up mass spectrometry has a significant limitation in that it currently struggles to reliably distinguish between differential regulation of a protein itself and its differential post-translational modification. The protein quantity quality score implemented in QuantUMS is not meant to address this problem. Nevertheless, we expect it to be a highly useful metric that can be used to improve downstream analysis by filtering out protein quantities significantly affected by MS-associated signal noise.

### Raw data processing

In-silico spectral libraries were generated from the sequence databases and refined on DIA data using DIA-NN 1.8.1. The PXD040205 data set (mixed species) was analysed with MS1 and MS2 mass accuracies fixed to 15 ppm and a scan window size of 6. The PXD022216 data set (leukaemia) was analysed with mass accuracies at 10 ppm, the window size was automatically inferred. Spectral libraries, DIA-NN pipeline files, logs and output reports have been deposited to the OSF repository https://osf.io/q8kfc/?view_only=5e77d3c62563468280fd09265583dbbd.

### Statistical analysis

Statistical analyses were performed in R 4.1. Differential expression analysis for the mixed species benchmark dataset was performed by applying the Welch‘s t-test to each log-transformed feature (precursor or protein). After ranking the features by their p-values, the effective false discovery rate (FDR) of each *E*.*coli* feature was calculated as the proportion of false (human) hits among all hits with a p-value lower than that of the considered *E*.*coli* feature. To calculate the differential expression detection rate, features were binned by their rank intensity and for each bin, the proportion of *E*.*coli* features with an effective FDR lower than a predefined threshold (0.05) was calculated. Differential abundance analysis for the CLL dataset was performed using the lmfit and eBayes functions of limma (v 3.50.3)^54^, fitting models on the log-transformed precursor or protein quantities separately for each of the patient characteristics in the supplied metadata.

## Data availability

The DIA-NN analysis reports and logs have been deposited to the OSF repository https://osf.io/q8kfc/?view_only=5e77d3c62563468280fd09265583dbbd. The raw proteomics data sets were downloaded from the repositories PXD040205 and PXD022216.

## Code availability

The scripts used to create the figures have been deposited to the OSF repository https://osf.io/q8kfc/?view_only=5e77d3c62563468280fd09265583dbbd. The DIA-NN version (1.8.2 beta 22) that implements QuantUMS is available by the same link. An open source release of QuantUMS, independent of the raw data processing software, is in preparation.

## Author contributions

Conception: V.D., F.K., J.G., L.R.S., algorithm design: F.K., software implementation: V.D., benchmarking: J.G., F.K., supervision: V.D.. All authors contributed to writing the manuscript.

## Acknowledgements

This work was supported by the German Ministry of Education and Research (BMBF), as part of the National Research Node “Mass spectrometry in Systems Medicine“ (MSCoreSys), under grant agreement 161L0221.

## Supplementary information

**Supplementary Figure S1.**
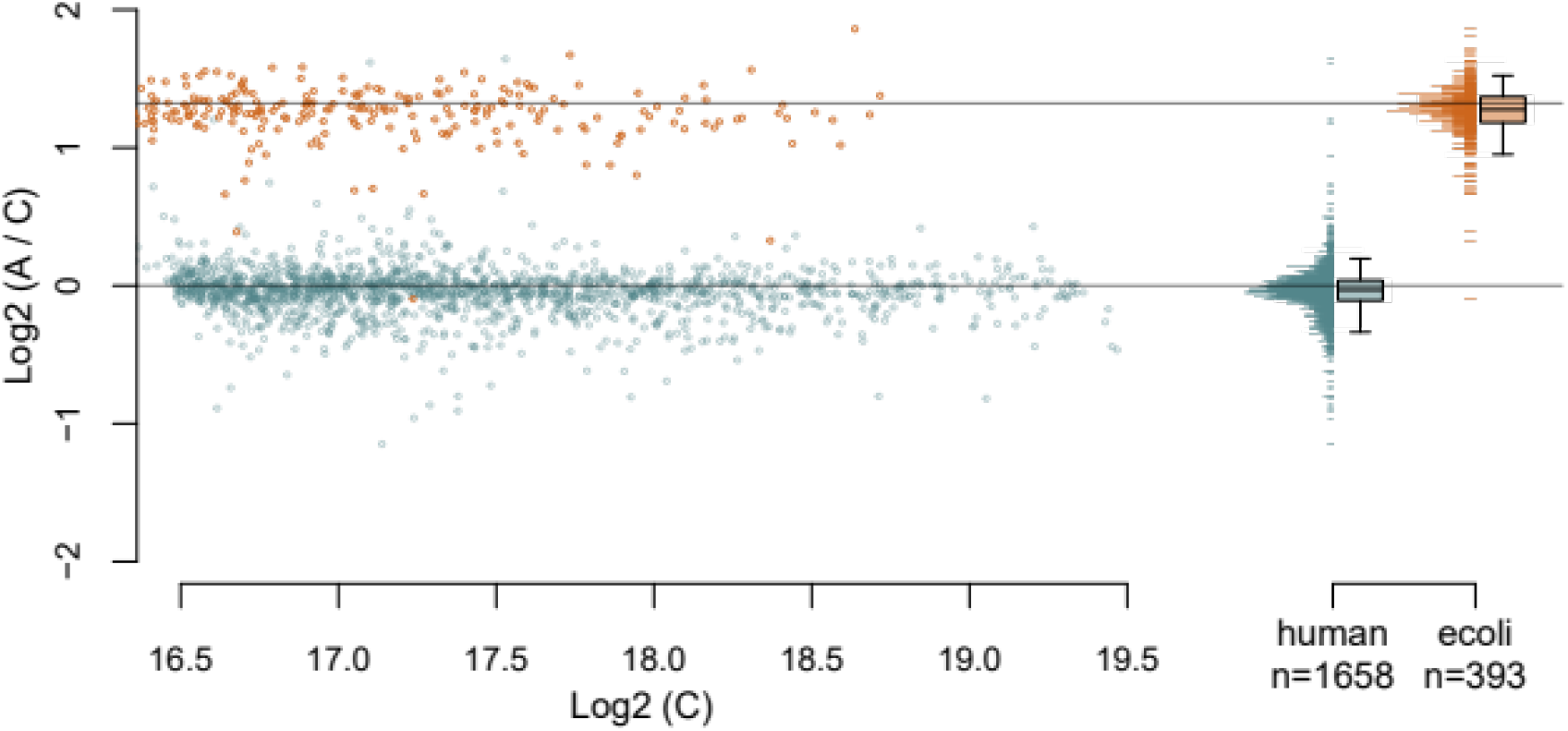
Validation of the *E*.*coli* dilution ratios. Mixtures A, B and C of human (K562) and *E*.*coli* tryptic digests with proportions A:B:C being 1:1:1 (human) and 50:33:20 (*E*.*coli*) were recorded^37^ using a 5-minute analytical flow gradient on timsTOF Pro and analysed with QuantUMS in high-accuracy mode. The DIA-NN output report was filtered for Lib.PG.Q.Value <= 0.01, PEP <= 0.01. Further, a stringent filter for MS1-based quantification was applied: Ms1.Profile.Corr > 0.95. The matrix of normalised MS1 precursor quantities was further filtered to only include precursors with median log2-quantities across samples A and C being between log2-transformed 0.85 and 0.99 quantiles of the Ms1.Area quantity reported by DIA-NN. Observed A:C ratios between thus filtered precursor MS1-level quantities were visualised. On the boxplots, boxes correspond to the interquartile range with the median indicated, whiskers extend to 5%-95% quantiles. Horizontal lines indicate the expected ratios for each species digest.

**Supplementary Figure S2.**
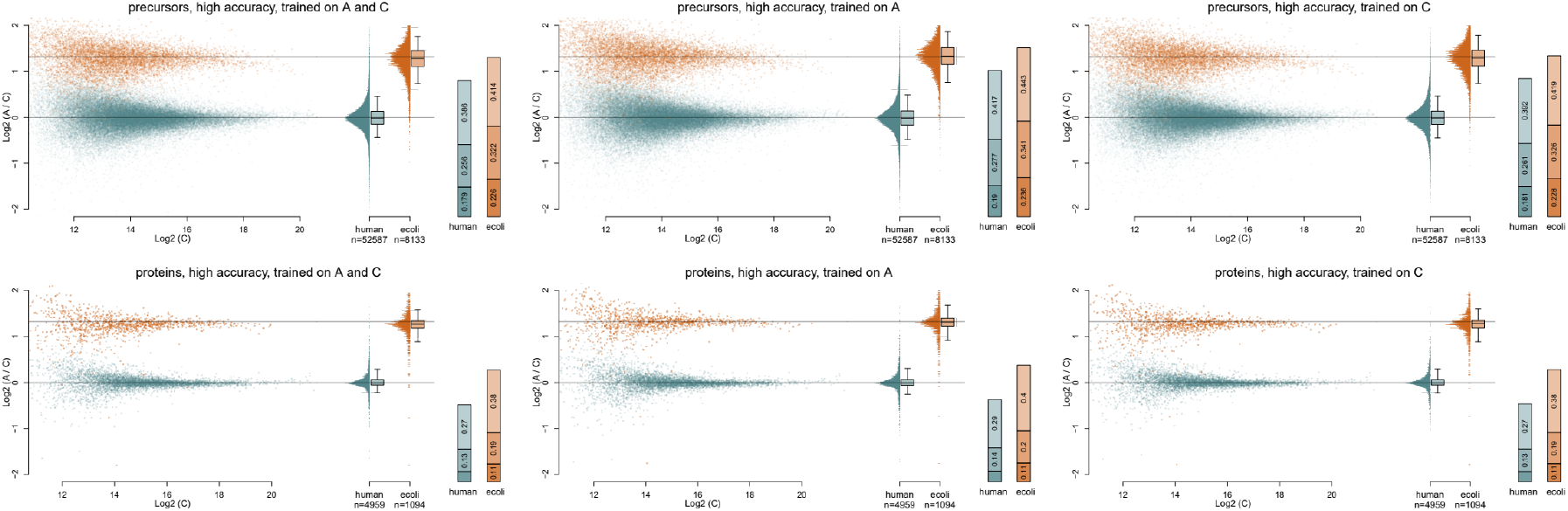
QuantUMS hyperparameter tuning performance does not depend on experimental design. Mixtures A, B and C of human (K562) and *E*.*coli* tryptic digests with proportions A:B:C being 1:1:1 (human) and 50:33:20 (*E*.*coli*) were recorded^37^ using a 5-minute analytical flow gradient on timsTOF Pro and analysed using DIA-NN. QuantUMS hyperparameters were trained on A+C, A and C subsets of the dataset separately, in high-accuracy mode, and then applied to the whole experiment. On the boxplots, boxes correspond to the interquartile range with the median indicated, whiskers extend to 5%-95% quantiles. Horizontal lines indicate the expected ratios for each species digest. Stacked bar plots indicate log2-standard deviations of A:C ratios within three intensity bins, with the lowest intensity bin at the top.

